# Cholesterol-Dependent Structure and Dynamics of Curved Lipid Vesicles Revealed by Dry MARTINI Simulations

**DOI:** 10.64898/2026.08.09.743827

**Authors:** Ehsan Khodadadi, Ehsaneh Khodadadi, Mahmoud Moradi

## Abstract

Cholesterol is a key regulator of membrane structure and dynamics, yet its effects on large curved vesicles under implicit-solvent coarse-grained conditions remain incompletely understood. Equilibrating large Dry MARTINI vesicles is challenging because transient membrane deformations can arise during the early stages of equilibration. Here, we developed a leaflet-specific restrained-equilibration protocol that preserves vesicle geometry while allowing local lipid relaxation. All restraints were removed before production simulations, and all reported results were obtained from unbiased trajectories. Using this protocol together with the Dry MARTINI force field and the TS2CG membrane builder, we simulated ∼50 nm DOPC vesicles containing 0–40 mol% cholesterol in three independent 20 µs production simulations for each membrane composition. Increasing cholesterol concentration produced a consistent structural reorganization of the membrane, characterized by increased membrane thickness and lipid-tail ordering, and decreased species-specific Voronoi area per lipid, lipid-tail interdigitation, solvent-accessible surface area, and vesicle shape anisotropy. Cholesterol flip-flop increased progressively with cholesterol concentration, whereas DOPC flip-flop exhibited a reproducible non-monotonic dependence with a maximum near 20 mol% cholesterol. Comparison with our previous explicit-solvent MARTINI simulations showed that the major cholesterol-dependent structural trends were preserved across both solvent representations, whereas species-specific lipid packing, lipid-tail interdigitation, and the absolute magnitude of lipid flip-flop remained sensitive to the solvent representation. Overall, Dry MARTINI combined with the restrained-equilibration protocol provides an efficient framework for studying large curved cholesterol-containing vesicles.

## 1. Introduction

Biological membranes are dynamic and compositionally heterogeneous structures whose physical properties emerge from the collective organization of lipids, sterols, and membrane-associated molecules. Membrane composition influences elasticity, permeability, curvature, lateral organization, and molecular transport, making lipid packing a fundamental determinant of membrane function. Understanding how composition regulates these properties is therefore important for both fundamental membrane biophysics and the rational design of lipid-based nanomaterials [1, 2, 3].

Molecular dynamics (MD) simulations provide a powerful approach for investigating membrane structure and dynamics at molecular resolution [4, 5]. However, atomistic simulations remain computationally demanding for large membrane systems and for processes occurring over microsecond-to-millisecond timescales [6]. Coarse-grained (CG) models address this limitation by reducing the number of interaction sites while retaining the principal physicochemical characteristics of the system, thereby extending the accessible spatial and temporal scales [7, 8, 9]. Among these models, the MARTINI force field has become one of the most widely used frameworks for membrane simulations because of its transferable coarse-grained parameterization and broad applicability to biomolecular systems [7, 10, 11, 8].

Despite the efficiency of conventional MARTINI, explicit solvent beads often account for most of the particles in large membrane simulations and therefore contribute substantially to the computational cost [12, 13]. Dry MARTINI was developed to reduce this burden by removing explicit water and reparameterizing the nonbonded interaction matrix to account for the omitted solvent degrees of freedom [14, 15]. The model reproduces a range of membrane properties reasonably well, including area per lipid, bilayer thickness, lipid-tail ordering, bending modulus, and membrane phase behavior. It has also been applied to large-scale processes such as membrane fusion and tether formation [14, 16]. Because the absence of explicit solvent substantially reduces the number of simulated particles, Dry MARTINI is particularly attractive for investigating large membrane assemblies over extended timescales. Nevertheless, the model is primarily qualitative, and some structural and dynamical properties can differ from those obtained using explicit-solvent MARTINI [14, 17].

Membrane curvature introduces an additional level of complexity by altering lipid packing, leaflet geometry, and the spatial organization of membrane components. The inner and outer leaflets of a vesicle have different surface areas and preferred radii, and these geometric differences can influence both local lipid organization and global vesicle shape. Direct self-assembly of large vesicles is computationally expensive, making the construction of realistic curved membrane models a significant practical challenge. The TS2CG (Triangulated Surface to Coarse-Grained) membrane builder addresses this limitation by generating CG membranes with user-defined geometry and composition from triangulated surfaces [18, 19]. This approach provides an efficient route for constructing large vesicles while preserving the intended membrane topology and leaflet composition.

Although geometry-based construction provides a suitable initial vesicle configuration, curved membranes require careful relaxation before production simulations. The original Dry MARTINI study also noted that deviations from planar bilayer properties introduced by vesicle curvature must be relaxed carefully before data collection [14]. In the present systems, preliminary unrestrained equilibration produced transient large-scale surface undulations and local membrane deformations during the early stages of relaxation. These changes did not represent the intended equilibrium sampling and complicated the preservation of the overall vesicle geometry during equilibration.

Collective-variable-based restraints provide a general strategy for controlling selected large-scale structural degrees of freedom while allowing local molecular relaxation [20, 21]. To address the equilibration challenge in the present systems, we developed a leaflet-specific restrained-equilibration protocol using radius-of-gyration collective variables. The inner and outer leaflets were restrained independently because membrane curvature gives rise to different characteristic radii for the two leaflets. These temporary restraints preserved the overall vesicle geometry while allowing local lipid rearrangement and packing relaxation. The restraint strength was introduced gradually during equilibration to avoid abrupt structural perturbations, and all restraints were removed before production simulations. Consequently, all structural and dynamical properties reported in this work were calculated from fully unrestrained production trajectories.

Dioleoylphosphatidylcholine (DOPC) is a widely used model phospholipid because it forms fluid bilayers under physiological conditions. Cholesterol is a major component of mammalian membranes and strongly influences lipid packing, membrane thickness, lateral organization, permeability, and molecular transport through its interactions with neighboring phospholipids. These effects are also important in liposomal drug-delivery systems, where cholesterol can alter membrane permeability, drug retention, and vesicle stability [22, 23]. Although the effects of cholesterol have been studied extensively in planar bilayers, its influence on large curved vesicles remains less well characterized, particularly under implicit-solvent coarse-grained conditions. In this study, we combined the Dry MARTINI force field, the TS2CG membrane builder, and a leaflet-specific restrained-equilibration protocol to investigate approximately 50 nm diameter DOPC vesicles containing 0, 10, 20, 30, and 40 mol% cholesterol. Three independent 20 µs production simulations were performed for each membrane composition. We evaluated the effects of cholesterol on membrane thickness, species-specific Voronoi packing descriptors, solvent-accessible surface area, lipid-tail ordering, radial interdigitation, transbilayer lipid flip-flop, and global vesicle shape anisotropy. By comparing these results with our previous explicit-solvent MARTINI simulations of the same DOPC/cholesterol vesicle system [24], we assess which cholesterol-dependent structural and dynamical trends are preserved across solvent representations and which remain sensitive to the choice of coarse-grained model.

## 2. Materials and Methods

### 2.1. Dry MARTINI Vesicle Construction

All simulations were performed using the Dry MARTINI CG force field, an implicit-solvent extension of the standard MARTINI model in which solvent-mediated effects are incorporated into the effective nonbonded interactions between CG particles [14]. By eliminating explicit solvent beads, Dry MARTINI substantially reduces the number of particles while reproducing many important structural and mechanical properties of lipid membranes. This improved computational efficiency makes the model well suited for microsecond-scale simulations of large membrane assemblies.

Initial vesicle configurations were generated using the TS2CG (Triangulated Surface to Coarse-Grained) membrane builder, version 2.0 [18, 19]. TS2CG constructs CG membranes from prescribed geometries using a two-stage workflow. First, the PLM (Pointillism) module discretizes the target surface into a point representation that preserves its local geometric properties, including surface area, coordinates, and curvature. The PCG module subsequently places lipid molecules onto this point distribution to generate an overlap-free membrane structure suitable for molecular dynamics simulations. For analytically defined geometries such as spherical vesicles, TS2CG generates the membrane directly without requiring an externally supplied triangulated mesh, thereby simplifying system preparation.

Compared with spontaneous self-assembly, this geometry-based approach allows membrane shape and lipid composition to be specified before simulation and produces near-equilibrium starting configurations, thereby reducing the extent of equilibration required before production simulations. Nevertheless, under the present Dry MARTINI simulation conditions, large curved vesicles required a dedicated restrained-equilibration protocol to preserve the overall vesicle geometry during the initial relaxation stage, as described in the following section.

Spherical vesicles with an initial diameter of approximately 50 nm were constructed containing 0, 10, 20, 30, and 40 mol% cholesterol, with DOPC comprising the remaining lipid fraction. This composition series was selected to systematically investigate the influence of cholesterol concentration on the structural and dynamical properties of large curved lipid vesicles.

Each vesicle was centered in a cubic simulation box with an edge length of 120 nm, and periodic boundary conditions were applied in all three dimensions. This box size provided a minimum separation of approximately 35 nm between the vesicle surface and its periodic images, which is substantially larger than the nonbonded interaction cutoffs used in this study (1.1–1.2 nm), thereby preventing spurious interactions between periodic images.

### 2.2. Simulation Protocol

All molecular dynamics simulations were performed using GROMACS 2024 [25]. Each system was first energy minimized using the steepest-descent algorithm for a maximum of 10 000 steps to remove unfavorable contacts introduced during vesicle construction. Electrostatic interactions were treated using the reaction-field method with a relative dielectric constant of *ɛ_r_* = 15 and a Coulomb cutoff of 1.2 nm. Lennard–Jones interactions were evaluated using a 1.2 nm cutoff together with the potential-shift modifier, and neighbor searching employed the Verlet cutoff scheme.

Following energy minimization, each vesicle was equilibrated using the stochastic dynamics (SD) integrator at 310 K with a 10 fs integration time step. Pressure coupling was not applied because the systems consisted of isolated vesicles simulated using the implicit-solvent Dry MARTINI model. Electrostatic interactions were treated using the reaction-field method with a relative dielectric constant of *ɛ_r_* = 15 and a Coulomb cutoff of 1.1 nm. Lennard–Jones interactions were evaluated using a 1.1 nm cutoff together with the potential-shift modifier, and neighbor searching employed the Verlet cutoff scheme.

Production simulations were subsequently performed using the stochastic dynamics integrator with a 4 fs integration time step at 310 K. The production simulation input additionally specified a velocity-rescaling thermostat with a coupling time constant of *τ_t_* = 4.0 ps. Pressure coupling was not applied. Electrostatic interactions were treated using the reaction-field method with a relative dielectric constant of *ɛ_r_* = 15 and a Coulomb cutoff of 1.1 nm, while Lennard–Jones interactions employed a 1.1 nm cutoff together with the potential-shift modifier. Neighbor searching employed the Verlet cutoff scheme. Center-of-mass translation was removed every 100 integration steps to prevent gradual accumulation of numerical drift during the long production simulations.

Each 20 µs production trajectory was generated as a series of sequentially restarted simulation segments, with each segment initialized from the final coordinates and velocities of the preceding run. This segmented execution was adopted to accommodate job scheduling limits on the high-performance computing system and did not interrupt the continuity of the molecular dynamics trajectories.

### 2.3. Radius-of-Gyration Restraints During Equilibration

Preliminary unrestrained simulations exhibited transient large-scale surface undulations and local membrane deformations during the initial stages of equilibration. To maintain the overall vesicle geometry while allowing local lipid relaxation, weak radius-of-gyration (*R_g_*) restraints were applied using the Collective Variables (Colvars) module [20, 8].

Because membrane curvature results in different equilibrium radii for the inner and outer leaflets, each leaflet was restrained independently. Two collective variables were defined using the membrane headgroup beads, including the PO4 beads of DOPC and the ROH beads of cholesterol, to monitor the radius of gyration of the outer and inner leaflets separately.

The outer leaflet was restrained within an *R_g_* range of 25.5–26.5 nm, while the inner leaflet was restrained between 23.5 and 24.5 nm, corresponding to their target equilibrium radii. Harmonic wall restraints were applied with a force constant that was gradually increased from 1.0 to 5.0 kJ mol*^−^*^1^ nm*^−^*^2^ over the first 100 000 simulation steps to avoid abrupt perturbations of the membrane structure during equilibration.

The restraints were applied only during the equilibration stage and were completely removed before production simulations. Consequently, all structural, dynamical, and mechanical properties reported in this work were calculated from fully unbiased production trajectories.

This restrained equilibration protocol follows the general collective-variable framework described by Fiorin *et al.* [20] and was designed to preserve the overall vesicle geometry while allowing local membrane relaxation before unbiased production simulations.

### 2.4. Production Simulations

Following equilibration and removal of all restraints, production simulations were performed according to the simulation protocol described in Section Simulation Protocol, using a 4 fs integration time step at 310 K under NVT conditions.

For each cholesterol concentration (0, 10, 20, 30, and 40 mol%), three independent production simulations of 20 µs were carried out using different random seeds, resulting in a total of 60 µs of sampling per composition and 300 µs across all systems. Unless otherwise stated, all analyses presented in this work were performed using these unbiased production trajectories.

### 2.5. Trajectory Analysis

Trajectory analyses were performed using custom Python scripts based on the MDAnalysis package [26, 27], together with NumPy [28] for numerical calculations and Matplotlib [29] for data visualization. Molecular structures and trajectories were visualized using Visual Molecular Dynamics (VMD) [30], while selected analyses, including solvent-accessible surface area (SASA), were performed using the GROMACS analysis tools [25]. Except where otherwise noted, all analyses were performed using the unbiased production trajectories following removal of the equilibration restraints.

For analyses that depended on vesicle geometry, the instantaneous center of geometry was calculated from the membrane headgroup beads. Radial coordinates were then used to account for the spherical geometry of the vesicles and to distinguish the inner and outer leaflets. Unless otherwise specified, the reported quantities represent global averages over the entire vesicle rather than measurements from selected local membrane regions.

The analyses included membrane thickness, species-specific Voronoi packing descriptors, SASA, the lipid-tail order parameter (*S_CD_*), radial lipid-tail interdigitation, lipid transbilayer exchange, and vesicle shape descriptors. Species-specific Voronoi packing descriptors were calculated by applying spherical Voronoi tessellation separately to the DOPC and cholesterol headgroup positions [31, 32]. These quantities were used to characterize the geometric packing associated with each lipid species and were not interpreted as a complete partitioning of the total leaflet surface area. SASA was calculated using the gmx sasa utility in GROMACS [25]. Vesicle morphology was characterized using the relative shape anisotropy (*κ*^2^) and normalized asphericity (*b*), calculated from the eigenvalues of the gyration tensor [33, 34]. The methodology for each analysis is described in the corresponding subsections below.

For each cholesterol composition, three independent 20 µs production simulations were analyzed. Composition-level values are reported as the mean across replicas, with the standard deviation among replicas used to quantify variability unless a figure caption specifies that block averaging within individual trajectories was used instead.

#### 2.5.1. Membrane Thickness

Membrane thickness was used to characterize the structural organization of the lipid bilayer and to assess the effect of cholesterol on membrane packing. Because the simulated systems are spherical vesicles, thickness was defined using a radial coordinate system rather than a planar membrane normal. For each trajectory frame, the instantaneous center of geometry of the vesicle was determined from all membrane headgroup beads, and lipids were assigned to the inner or outer leaflet according to their radial positions.

The instantaneous membrane thickness was calculated as the difference between the average radial positions of the outer and inner leaflet headgroups,

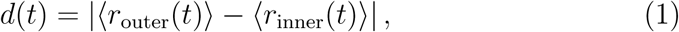

where *r* is the distance of each lipid headgroup from the instantaneous vesicle center and ⟨·⟩ denotes the average over all headgroups within the corresponding leaflet. The equilibrium membrane thickness for each system was obtained by averaging the instantaneous thickness over the production trajectory.

This radial definition is well suited for highly curved membrane systems because it naturally accounts for the spherical geometry of the vesicle without requiring the definition of a local membrane normal. Membrane thickness was calculated independently for each simulation replica, and the reported values correspond to the mean and standard deviation of the three independent production simulations for each cholesterol concentration.

This approach provides a robust global measure of membrane organization and enables quantitative comparison of cholesterol-dependent changes in bilayer thickness across curved vesicles [35, 31].

#### 2.5.2. Radial Tail Interdigitation

Lipid-tail interdigitation was quantified using a radial adaptation of the count-based approach previously developed for planar and spherical membranes [24], which is based on established bead-overlap methods for lipid bilayers [36, 37, 38]. Because spherical vesicles lack a single global membrane normal, the analysis was expressed in terms of the radial distance from the instantaneous center of geometry of the vesicle rather than a planar membrane normal.

For each trajectory frame, the instantaneous center of geometry was calculated from all membrane headgroup beads, and lipids were assigned to the inner or outer leaflet according to the radial position of their headgroups. The instantaneous bilayer midplane was then defined as the midpoint between the average radial positions of the two leaflets. The radial position of each DOPC tail bead was calculated as

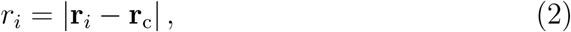

where **r***_i_* is the position vector of tail bead *i* and **r**_c_ is the instantaneous center of geometry of the vesicle. A tail bead was classified as interdigitating if its radial position crossed the instantaneous bilayer midplane into the opposite leaflet. The interdigitation value was then calculated as the average number of crossing tail beads per lipid for each leaflet at each trajectory frame.

Expressing the analysis in radial coordinates makes the method appropriate for highly curved vesicles while preserving the same count-based definition used in our previous explicit-solvent study [24]. This consistent definition enables direct comparison of cholesterol-dependent lipid-tail interdigitation between the explicit- and implicit-solvent MARTINI models.

#### 2.5.3. Species-Specific Voronoi Packing Descriptors

Species-specific lipid packing descriptors were calculated using spherical Voronoi tessellation, which provides a geometry-based measure of local lipid packing without assuming a predefined lattice arrangement [39, 40]. Because the simulated systems are closed spherical vesicles, the analysis was performed directly on the vesicle surface rather than using the planar Voronoi approach commonly applied to flat membranes.

For each trajectory frame, the instantaneous center of geometry of the vesicle was determined from all membrane headgroup beads. The phosphate (PO4) bead of DOPC and the hydroxyl (ROH) bead of cholesterol were used as representative headgroup positions according to the MARTINI mapping scheme [41]. Lipids were assigned to the inner or outer leaflet based on the radial position of their headgroups using the same clustering procedure applied throughout this work to distinguish the two leaflets.

Within each leaflet, the headgroup coordinates of DOPC and cholesterol were analyzed separately. The headgroup positions were projected onto a unit sphere centered at the vesicle center while preserving their angular distribution. A spherical Voronoi tessellation was then constructed, and the area of each Voronoi cell was calculated on the unit sphere using the spherical excess method. The corresponding physical surface area was obtained by scaling each Voronoi cell by the square of the mean leaflet radius for the corresponding lipid species,

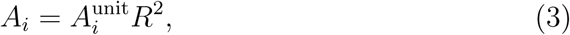

where *A_i_*^unit^ is the Voronoi cell area on the unit sphere and *R* is the mean radial distance of the corresponding lipid species within the leaflet for that trajectory frame. The average area was obtained by averaging the Voronoi cell areas over all molecules of the same species within each leaflet. Population-weighted averages of the inner and outer leaflets were then used to obtain the overall average area for each lipid species.

Because DOPC and cholesterol were tessellated independently rather than within a mixed-species Voronoi diagram, the calculated areas characterize the local geometric packing associated with each lipid species separately. The reported values should therefore be interpreted as species-specific packing descriptors rather than as a complete partitioning of the total leaflet surface area.

#### 2.5.4. Solvent-Accessible Surface Area

The SASA was calculated to evaluate changes in membrane surface exposure and lipid packing as a function of cholesterol concentration. Although the Dry MARTINI model employs an implicit-solvent representation, SASA remains a useful geometric descriptor because it quantifies the molecular surface accessible to a probe of fixed radius, independent of the solvent model. In this study, SASA was therefore used as a structural measure of membrane packing rather than as a direct measure of solvent accessibility or membrane permeability.

SASA calculations were performed using the gmx sasa module in GRO-MACS [25], which implements the Shrake–Rupley rolling-probe algorithm [42]. The same probe radius was used for all simulations to ensure consistent comparisons across membrane compositions.

The solvent-accessible surface area of the complete membrane and the species-specific SASA of DOPC and cholesterol were analyzed independently. The species-specific SASA values were used as geometric descriptors of the surface associated with each lipid species and were not intended to represent additive contributions to the total membrane SASA. To account for the different numbers of DOPC and cholesterol molecules in each system, species-specific SASA values were normalized on a per-lipid basis before averaging. Because the simulated systems were closed spherical vesicles, SASA was evaluated over the entire vesicle surface.

Species-normalized SASA values reflect both membrane organization and changes in lipid composition as cholesterol progressively replaces DOPC. Consequently, species-specific SASA should be interpreted as a compositional structural descriptor rather than a direct measure of the local surface exposure of individual lipid molecules. Together with membrane thickness, the species-specific Voronoi packing descriptors, lipid-tail interdigitation, and the lipid-tail order parameter (*S_CD_*), SASA provides a complementary measure of cholesterol-dependent structural changes in Dry MARTINI vesicles.

#### 2.5.5. Lipid-Tail Order Parameter

The orientational ordering of the DOPC acyl chains was quantified using a CG lipid-tail order parameter, denoted *S_CD_*, calculated from the second Legendre polynomial of bond vectors connecting consecutive MARTINI tail beads. Because the MARTINI force field does not explicitly represent C–H bonds, this quantity should be regarded as a CG analogue of the deuterium order parameter and was used here to compare relative lipid-tail ordering across membrane compositions.

The order parameter was calculated as

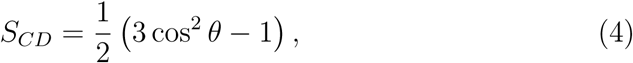

where *θ* is the angle between the bond vector connecting two consecutive CG tail beads and the local membrane normal. Values approaching 1 indicate preferential alignment of the bond vector with the membrane normal, whereas lower values indicate greater orientational disorder. This expression corresponds to the second Legendre polynomial and is widely used to characterize lipid-chain orientational ordering [43, 44].

For each trajectory frame, the instantaneous center of geometry of the vesicle was determined from all membrane headgroup beads. The local membrane normal for each DOPC molecule was defined as the radial vector connecting the vesicle center to the phosphate (PO4) bead, thereby accounting for the spherical membrane geometry. Bond vectors were calculated between consecutive beads along both DOPC acyl chains: C1A–D2A, D2A–C3A, and C3A–C4A for the sn-1 chain, and C1B–D2B, D2B–C3B, and C3B–C4B for the sn-2 chain. The order parameter was calculated for each bond segment and averaged over the DOPC molecules and production trajectory. Segment-level values were subsequently combined to obtain a representative order parameter for each acyl chain at each cholesterol concentration.

Higher *S_CD_* values indicate greater orientational alignment of the lipid-tail bond vectors with the local membrane normal and, therefore, a higher degree of lipid-tail ordering. The calculated values were used to quantify the effect of cholesterol concentration on DOPC acyl-chain ordering in the Dry MARTINI vesicles.

#### 2.5.6. Lipid Transbilayer Exchange

Lipid transbilayer exchange (flip-flop) was analyzed to quantify the translocation of DOPC and cholesterol molecules between the inner and outer leaflets of the spherical vesicles. For each trajectory frame, the instantaneous center of geometry of the vesicle was determined from all membrane headgroup beads, and the radial distance of each headgroup from the vesicle center was calculated. Lipids were first assigned to the inner or outer leaflet using the same clustering procedure applied throughout this work, and the instantaneous bilayer midplane was defined as the midpoint between the average radial positions of the two leaflets. This radial approach naturally accounts for the spherical geometry of the vesicle.

Each lipid was subsequently monitored according to the radial position of its headgroup relative to the instantaneous bilayer midplane. To avoid spurious leaflet transitions caused by thermal fluctuations near the membrane center, a hysteresis region was applied around the midplane, and a leaflet reassignment was recorded only after the headgroup had fully entered the opposite leaflet. A translocation event was classified as a flip-flop only if the lipid remained continuously within the new leaflet for at least 50 ns, thereby excluding transient midplane crossings.

Confirmed inner-to-outer and outer-to-inner translocation events were counted throughout the production trajectories. Flip-flop frequencies were calculated as the number of confirmed translocation events per lipid per microsecond and averaged over the three independent production simulations for each cholesterol concentration.

Because lipid flip-flop rates in coarse-grained simulations depend on the underlying force field, the calculated flip-flop frequencies were therefore used to compare the relative effects of cholesterol concentration within the Dry MARTINI model rather than to estimate experimental flip-flop rates directly.

#### 2.5.7. Quantification of Vesicle Shape Fluctuations

Global vesicle morphology was characterized using two complementary shape descriptors: the relative shape anisotropy (*κ*^2^) and the normalized asphericity (*b*), using the gyration-tensor formalism originally developed by Rudnick and Gaspari [33] and Aronovitz and Nelson [34], following the implementation described by Arkın and Janke [45]. Gyration-tensor-based descriptors have been widely applied to characterize the morphology of nanoscale systems in molecular dynamics simulations [46]. These descriptors were calculated for all cholesterol concentrations (0, 10, 20, 30, and 40 mol%) using the membrane headgroup beads, including the phosphate (PO4) beads of DOPC and the hydroxyl (ROH) beads of cholesterol.

For each trajectory frame, the instantaneous center of geometry of the selected headgroup beads was calculated, and the gyration tensor was constructed as

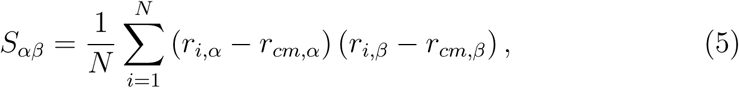

where *N* is the number of selected headgroup beads, *r_i_* is the position vector of bead *i*, *r_cm_* is the center of geometry of the vesicle, and *α, β* ∈ {*x, y, z*}.

Diagonalization of the gyration tensor yields the three principal eigenvalues,

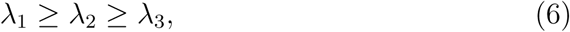

which describe the spatial distribution of the membrane along the three principal axes.

The relative shape anisotropy was calculated as

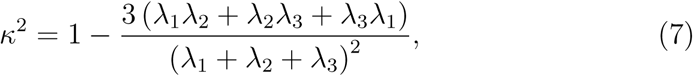

where *κ*^2^ = 0 corresponds to a perfectly isotropic (spherical) vesicle, and increasing values indicate progressively greater deviations from spherical symmetry.

The normalized asphericity was calculated as

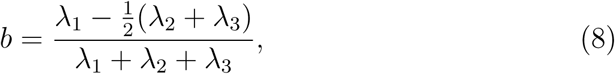

which provides a complementary measure of vesicle elongation based on the relative extension of the principal axes.

Both descriptors were calculated for every trajectory frame throughout the production simulations, allowing the temporal evolution of vesicle shape to be monitored for each cholesterol concentration. Because the normalized asphericity showed the same qualitative trend with cholesterol concentration as the relative shape anisotropy, only *κ*^2^ is presented in the Results for clarity.

## 3. Results and Discussion

### 3.1. Effect of Cholesterol on Membrane Thickness

Membrane thickness is a fundamental structural property of lipid bilayers that reflects changes in lipid packing and membrane organization. As shown in Figure 1(Left), membrane thickness increases progressively with cholesterol concentration in all three independent simulations. The average thickness increases from approximately 39.5–39.9 Å for the cholesterol-free vesicle to approximately 42.5–42.7 Å at 40 mol% cholesterol, corresponding to an overall increase of about 3 Å. Most of this increase occurs between 0 and 20 mol% cholesterol, whereas only a modest additional increase is observed between 20 and 40 mol%, suggesting that membrane thickness approaches a plateau at higher cholesterol concentrations.

**Figure 1:**
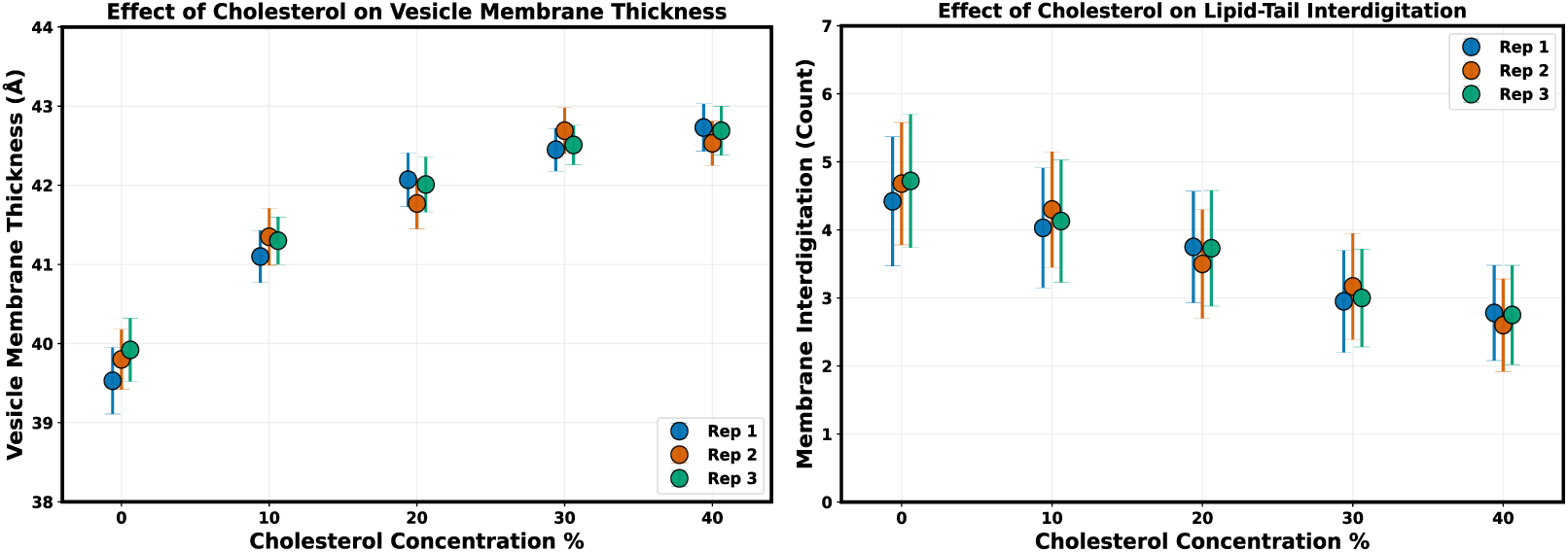
Effect of cholesterol concentration on membrane thickness and lipidtail interdigitation in Dry MARTINI vesicles. (Left) Equilibrium membrane thickness of approximately 50 nm DOPC vesicles containing 0–40 mol% cholesterol. Membrane thickness increases progressively with cholesterol concentration, reflecting cholesterolinduced membrane condensation and lipid-tail ordering. (Right) Lipid-tail interdigitation measured using the radial count-based method. Interdigitation decreases with increasing cholesterol concentration, indicating reduced overlap of hydrocarbon chains between opposing leaflets. Each symbol represents one of three independent 20 µs production simulations. Error bars represent the standard deviation obtained from block averaging within each simulation.

This behavior is consistent with the well-established condensing effect of cholesterol. By restricting the conformational flexibility of neighboring DOPC acyl chains, cholesterol promotes more extended lipid conformations and tighter molecular packing, leading to an increase in bilayer thickness along the membrane normal [47, 48, 49]. The close agreement among the three independent simulations demonstrates that the observed cholesterol-dependent thickening is highly reproducible.

The same trend was observed in our previous explicit-solvent MARTINI simulations of curved DOPC/cholesterol vesicles [24], suggesting that the cholesterol-induced increase in membrane thickness is preserved across both explicit- and implicit-solvent MARTINI representations.

### 3.2. Effect of Cholesterol on Lipid-Tail Interdigitation

Lipid-tail interdigitation measures the extent to which hydrocarbon chains from opposing leaflets penetrate across the bilayer midplane. As shown in Figure 1(Right), interdigitation decreases steadily with increasing cholesterol concentration in all three independent simulations. The average value decreases from approximately 4.4–4.7 crossing tail beads per lipid at 0 mol% cholesterol to approximately 2.6–2.8 at 40 mol% cholesterol, corresponding to an overall reduction of about 40%.

This decrease reflects the ordering effect of cholesterol on lipid tails. As cholesterol promotes tighter lipid packing and more extended acyl-chain conformations, penetration of hydrocarbon chains across the bilayer midplane becomes progressively less favorable, resulting in reduced interleaflet overlap. Unlike membrane thickness, which begins to level off above 20 mol% cholesterol, interdigitation continues to decrease throughout the concentration range investigated, indicating that interleaflet packing remains responsive to cholesterol even after membrane thickness begins to approach a plateau.

The contrasting concentration dependence of these two structural properties indicates that membrane thickness approaches a near-equilibrium value at intermediate cholesterol concentrations, whereas interleaflet packing continues to respond over the full cholesterol range investigated.

Compared with our previous explicit-solvent MARTINI simulations [24], the reduction in interdigitation is more pronounced in the Dry MARTINI model. This observation suggests that interdigitation is more sensitive to the underlying solvent representation than membrane thickness, although both models predict the same overall ordering effect of cholesterol.

Taken together, the increase in membrane thickness and the simultaneous decrease in lipid-tail interdigitation provide complementary structural descriptors of the same cholesterol-induced membrane reorganization. Whereas membrane thickness reflects extension of the lipid tails along the membrane normal, interdigitation measures the extent of overlap between the two opposing leaflets. The opposite trends observed for these two descriptors indicate that tighter lipid packing is accompanied by reduced interleaflet penetration and the formation of a more ordered bilayer architecture.

### 3.3. Cholesterol-Dependent Area per Lipid

The species-resolved average area per lipid obtained from the spherical Voronoi analysis is shown in Figure 2. Panels A and B show the average Voronoi area associated with cholesterol and DOPC molecules, respectively, as a function of cholesterol concentration.

**Figure 2:**
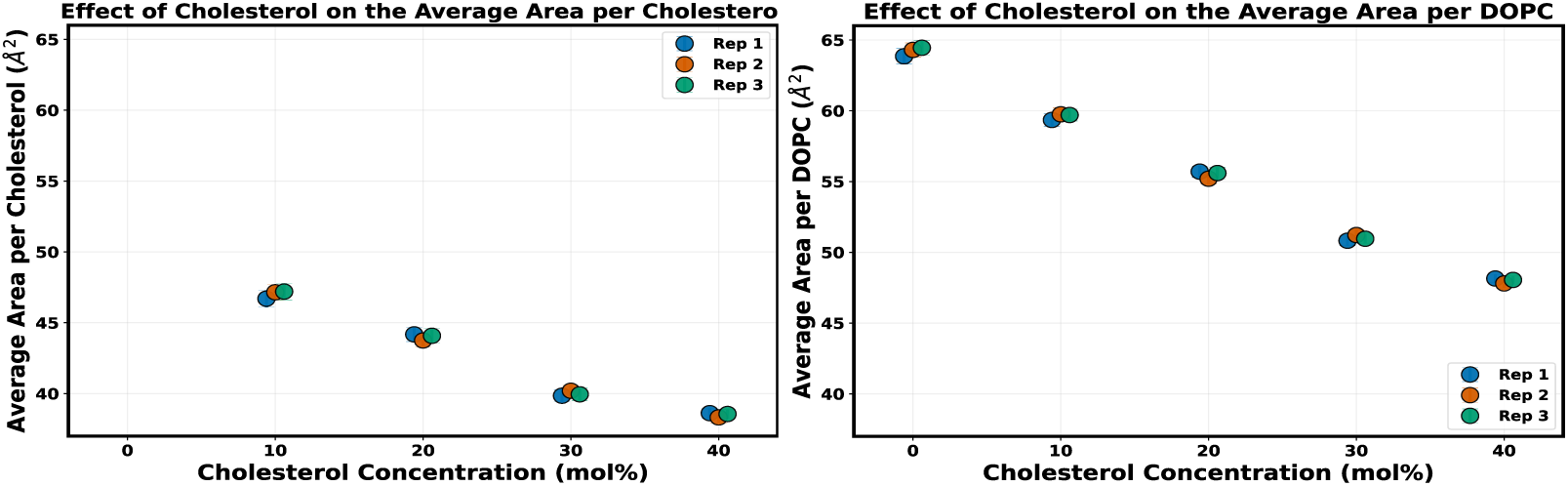
Effect of cholesterol concentration on the species-resolved average area per lipid in Dry MARTINI vesicles. (Left) Average Voronoi area per cholesterol molecule for approximately 50 nm DOPC vesicles containing 10–40 mol% cholesterol. (Right) Average Voronoi area per DOPC molecule for vesicles containing 0–40 mol% cholesterol. Each symbol represents one of three independent 20 µs production simulations. Both cholesterol and DOPC exhibit progressively smaller species-specific Voronoi areas with increasing cholesterol concentration, consistent with progressive membrane condensation. Error bars represent the standard deviation obtained from block averaging within each simulation.

#### 3.3.1. Effect of Cholesterol on the Average Area per Cholesterol Molecule

Because the cholesterol-free system contains no cholesterol molecules, the cholesterol-specific area is reported only for systems containing 10–40 mol% cholesterol. As shown in Figure 2(Left), the average area associated with cholesterol decreases steadily as the cholesterol content of the membrane increases. The average area per cholesterol molecule decreases from approximately 47 Å^2^ at 10 mol% cholesterol to about 38.5 Å^2^ at 40 mol%, corresponding to an overall reduction of nearly 20%. The close agreement among the three independent simulations demonstrates that this concentration-dependent trend is highly reproducible.

The progressive reduction in the cholesterol-specific Voronoi area indicates that the effective lateral footprint occupied by cholesterol becomes smaller as cholesterol concentration increases, consistent with progressively tighter lipid packing within the membrane.

#### 3.3.2. Effect of Cholesterol on the Average Area per DOPC Molecule

A similar trend is observed for DOPC (Figure 2(Right)). The average area per DOPC molecule decreases progressively from approximately 64 Å^2^ in the cholesterol-free membrane to about 48 Å^2^ at 40 mol% cholesterol. As with cholesterol, the three independent simulations show excellent agreement, demonstrating the robustness of the observed concentration dependence.

By restricting the conformational flexibility of neighboring acyl chains, cholesterol promotes tighter packing of DOPC molecules, reducing the average lateral area occupied by each phospholipid. This trend is consistent with the other structural changes observed throughout this study. As membrane thickness increases and lipid tails become more ordered, each lipid occupies a smaller lateral area, accompanied by reduced interdigitation between opposing leaflets.

Taken together, the reductions in both cholesterol and DOPC Voronoi areas provide complementary evidence that increasing cholesterol drives the membrane toward a more compact organization. This behavior differs from our previous explicit-solvent MARTINI simulations, in which the species-resolved Voronoi areas associated with cholesterol and DOPC exhibited opposite concentration-dependent trends [24]. In contrast, the Dry MARTINI model predicts a parallel decrease in the Voronoi area associated with both lipid species. This difference suggests that species-specific lipid packing is more sensitive to the underlying solvent representation than the overall cholesterol-induced membrane condensation process. Nevertheless, the global structural response to increasing cholesterol remains consistent across both models, including increased membrane thickness, enhanced lipid-tail ordering, reduced interdigitation, decreased solvent-accessible surface area, and an overall increase in membrane packing.

### 3.4. Effect of Cholesterol on Solvent-Accessible Surface Area

SASA provides a quantitative measure of the membrane surface exposed to the surrounding environment and is commonly used to assess changes in membrane packing and molecular exposure. Figure 3 shows the average SASA per lipid for the entire membrane as a function of cholesterol concentration.

**Figure 3:**
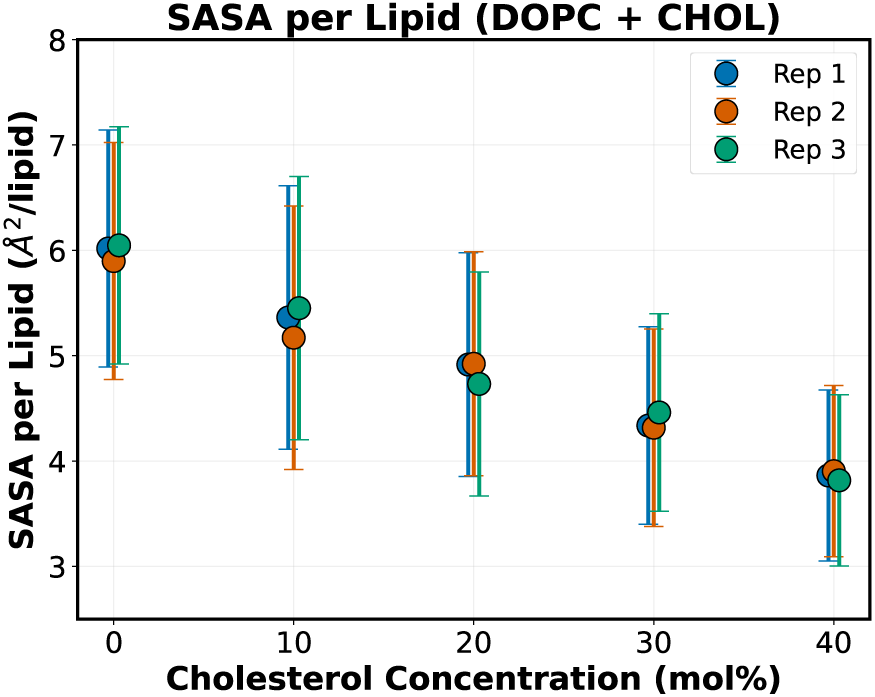
Effect of cholesterol concentration on the total SASA per lipid in Dry MARTINI vesicles. The average SASA per lipid (DOPC + CHOL) decreases progressively with increasing cholesterol concentration, indicating reduced membrane surface exposure as cholesterol promotes tighter membrane packing. Each symbol represents one of three independent 20 µs production simulations. Error bars represent the standard deviation obtained from block averaging within each simulation.

#### 3.4.1. Effect of Cholesterol on the Total Membrane SASA

As shown in Figure 3, the average SASA per lipid decreases progressively with increasing cholesterol concentration. The average SASA decreases from approximately 6.0 Å^2^/lipid in the cholesterol-free membrane to about 3.8 Å^2^/lipid at 40 mol% cholesterol, corresponding to an overall reduction of nearly 35%. The close agreement among the three independent simulations demonstrates that this concentration-dependent trend is highly reproducible. The reduction in total SASA indicates that progressively less membrane surface remains accessible to the surrounding environment as cholesterol concentration increases. This trend is fully consistent with the structural changes described in the preceding sections. Reduced solvent exposure accompanies the decrease in Voronoi area per lipid and lipid-tail interdigitation, together with the increase in membrane thickness and lipid-tail ordering, providing independent evidence that cholesterol promotes progressively tighter membrane packing.

#### 3.4.2. Species-Resolved Solvent-Accessible Surface Area

The species-resolved SASA of cholesterol and DOPC is shown in Figure 4. Because the cholesterol-free system contains no cholesterol molecules, the cholesterol-specific SASA is reported only for systems containing 10–40 mol% cholesterol.

**Figure 4:**
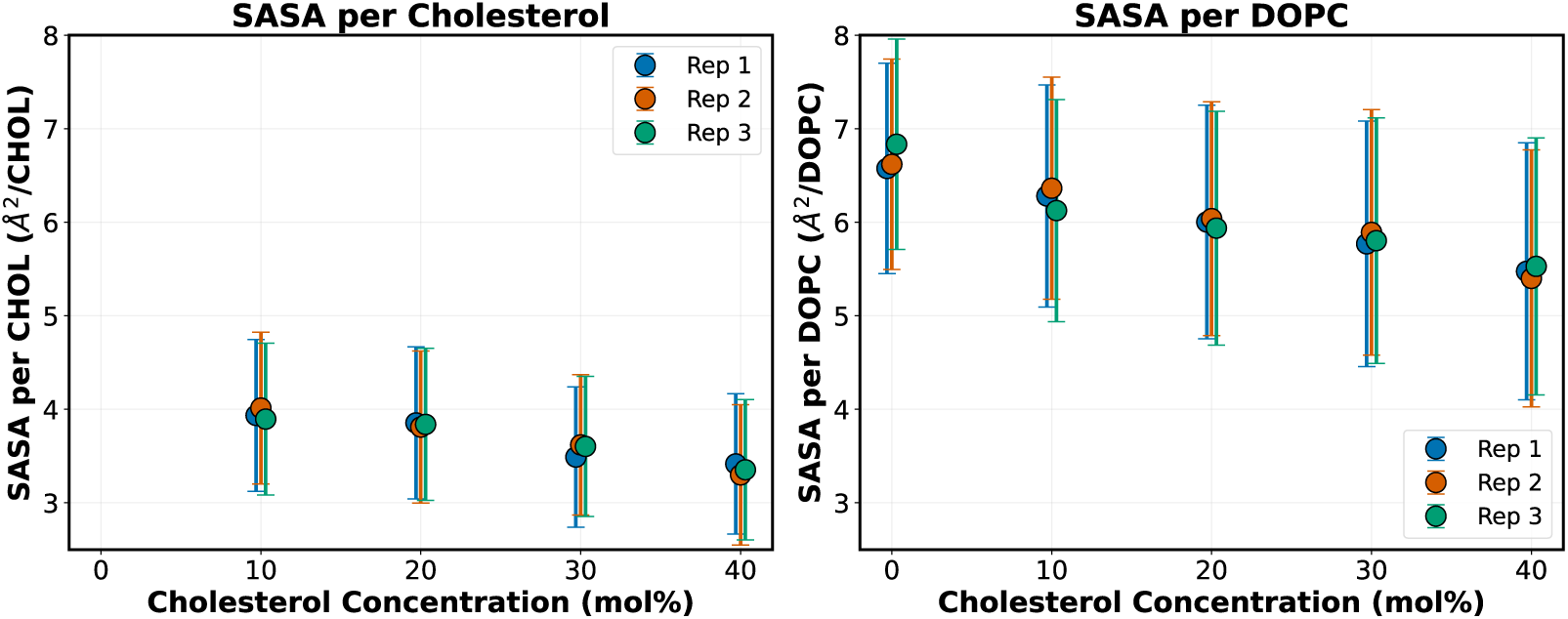
Species-resolved SASA in Dry MARTINI vesicles. (Left) Average SASA per cholesterol molecule for approximately 50 nm DOPC vesicles containing 10– 40 mol% cholesterol. (Right) Average SASA per DOPC molecule for vesicles containing 0–40 mol% cholesterol. Each symbol represents one of three independent 20 µs production simulations. Both cholesterol and DOPC exhibit progressively lower average SASA with increasing cholesterol concentration, indicating reduced solvent exposure of both lipid species. Error bars represent the standard deviation obtained from block averaging within each simulation.

As shown in Figure 4(Left), the average SASA per cholesterol molecule decreases from approximately 4.0 Å^2^/molecule at 10 mol% cholesterol to about 3.3 Å^2^/molecule at 40 mol% cholesterol.

A similar trend is observed for DOPC (Figure 4(Right)). The average SASA per DOPC molecule decreases from approximately 6.6 Å^2^/molecule in the cholesterol-free membrane to about 5.5 Å^2^/molecule at 40 mol% cholesterol. In both cases, the three independent simulations show excellent agreement across the entire composition range, confirming that this behavior is consistent across independent trajectories.

The parallel reduction in SASA for both lipid species indicates that increasing cholesterol reduces the solvent exposure of both sterol and phospholipid molecules. The decrease in cholesterol SASA reflects progressively closer sterol packing, whereas the reduction in DOPC SASA indicates that phospholipid molecules also become less exposed as the membrane becomes more tightly packed.

Because the species-resolved SASA values are calculated independently for each lipid species, they should be interpreted as complementary descriptors of the average solvent exposure associated with cholesterol and DOPC rather than as additive contributions to the total membrane SASA.

Taken together, the total and species-resolved SASA analyses provide a consistent picture of cholesterol-induced membrane remodeling. Increasing cholesterol reduces both the overall membrane surface exposure and the average accessible surface associated with each lipid species. These observations complement the reductions in Voronoi area per lipid and lipid-tail interdigitation, together with the increases in membrane thickness and lipid-tail ordering, demonstrating that cholesterol progressively drives the membrane toward a more ordered and tightly packed structure.

### 3.5. Effect of Cholesterol on Lipid-Tail Ordering

The orientational ordering of the DOPC acyl chains was quantified using the lipid-tail order parameter (*S_CD_*). As shown in Figure 5, increasing cholesterol concentration produces a progressive increase in *S_CD_* for both the sn-1 and sn-2 acyl chains. The close agreement among the three independent simulations demonstrates that the observed concentration-dependent trends are highly reproducible.

**Figure 5:**
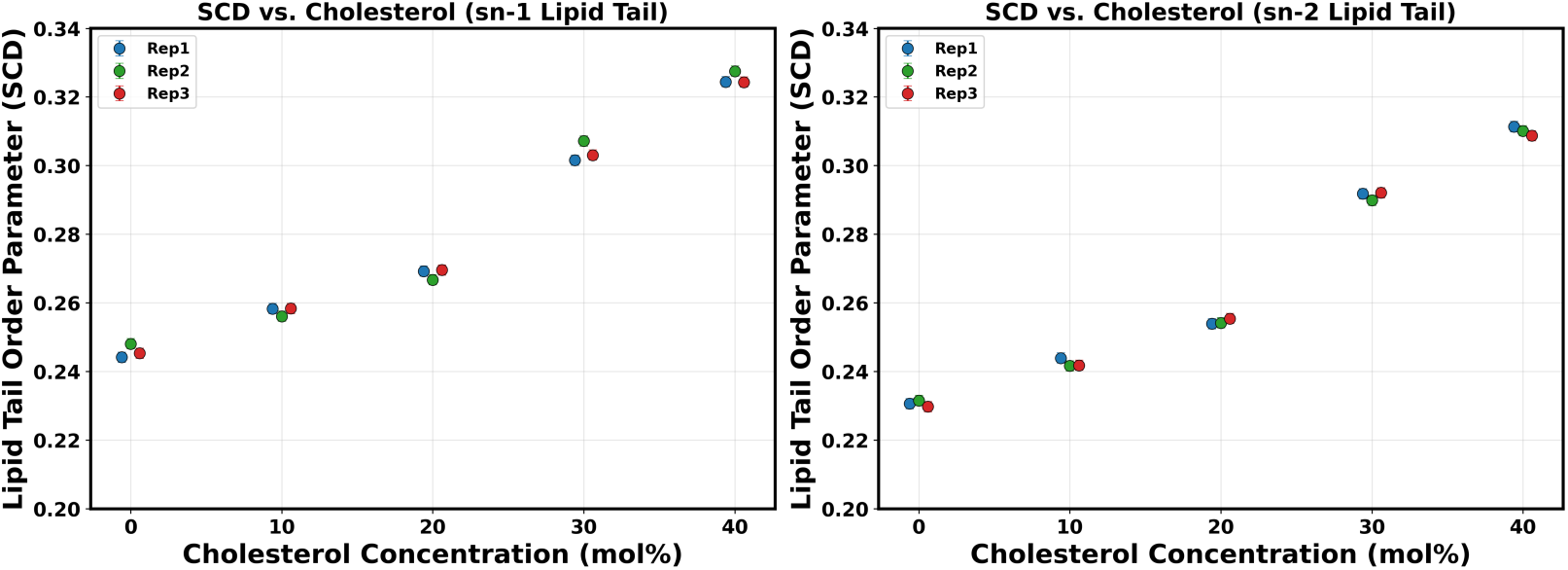
**Effect of cholesterol concentration on the lipid-tail order parameter (***S_CD_***) of DOPC acyl chains in Dry MARTINI vesicles.** (Left) Average lipid-tail order parameter of the sn-1 acyl chain as a function of cholesterol concentration. (Right) Average lipid-tail order parameter of the sn-2 acyl chain as a function of cholesterol concentration. Both acyl chains exhibit a progressive increase in *S_CD_* with increasing cholesterol concentration, indicating enhanced lipid-tail ordering. The sn-1 chain maintains slightly higher *S_CD_* values than the sn-2 chain across all membrane compositions. Each symbol represents one of three independent 20 µs production simulations. Error bars represent the standard deviation obtained from block averaging within each simulation.

For the sn-1 chain (Figure 5(Left)), the average *S_CD_* increases from approximately 0.245 in the cholesterol-free membrane to about 0.325 at 40 mol% cholesterol. Similarly, the average *S_CD_* of the sn-2 chain (Figure 5(Right)) increases from approximately 0.231 to about 0.311 over the same concentration range. For both chains, the increase is relatively gradual between 0 and 20 mol% cholesterol, becomes more pronounced between 20 and 30 mol%, and continues between 30 and 40 mol%. Thus, no clear plateau in lipid-tail ordering is observed within the concentration range investigated.

The increase in *S_CD_* is consistent with the well-established ordering effect of cholesterol on phospholipid membranes. The rigid sterol ring restricts the conformational flexibility of neighboring acyl chains and favors more extended chain conformations aligned more closely with the membrane normal [43, 50, 51]. This ordering response is consistent with the accompanying increase in membrane thickness and the reductions in lipid-tail interdigitation, area per lipid, and solvent-accessible surface area. Together, these structural changes indicate that increasing cholesterol promotes a more ordered and tightly packed DOPC membrane.

Although both acyl chains respond similarly to increasing cholesterol concentration, the sn-1 chain consistently exhibits slightly higher *S_CD_* values than the sn-2 chain across all membrane compositions. This difference has previously been reported for phosphatidylcholine bilayers and may reflect the distinct conformational and molecular environments of the two acyl chains within the phospholipid molecule [50]. Nevertheless, the nearly parallel concentration-dependent increases observed for both chains indicate that cholesterol enhances ordering throughout both acyl chains rather than preferentially affecting only one chain.

### 3.6. Flip-Flop Dynamics

Lipid flip-flop was analyzed to quantify transbilayer migration between the inner and outer leaflets of the spherical Dry MARTINI vesicles. Flip-flop frequencies were calculated separately for outer-to-inner and inner-to-outer translocation events for both cholesterol and DOPC.

#### 3.6.1. Cholesterol Flip-Flop

The cholesterol flip-flop frequencies are shown in Figure 6. Both outer-to-inner (Figure 6(Left)) and inner-to-outer (Figure 6(Right)) translocation frequencies increase progressively with cholesterol concentration. At 10 mol% cholesterol, only a small number of transbilayer exchange events are observed, whereas substantially higher flip-flop frequencies occur at 30 and 40 mol% cholesterol. The two translocation directions closely follow one another across the entire concentration range, indicating that cholesterol exchange remains approximately balanced between the inner and outer leaflets.

**Figure 6:**
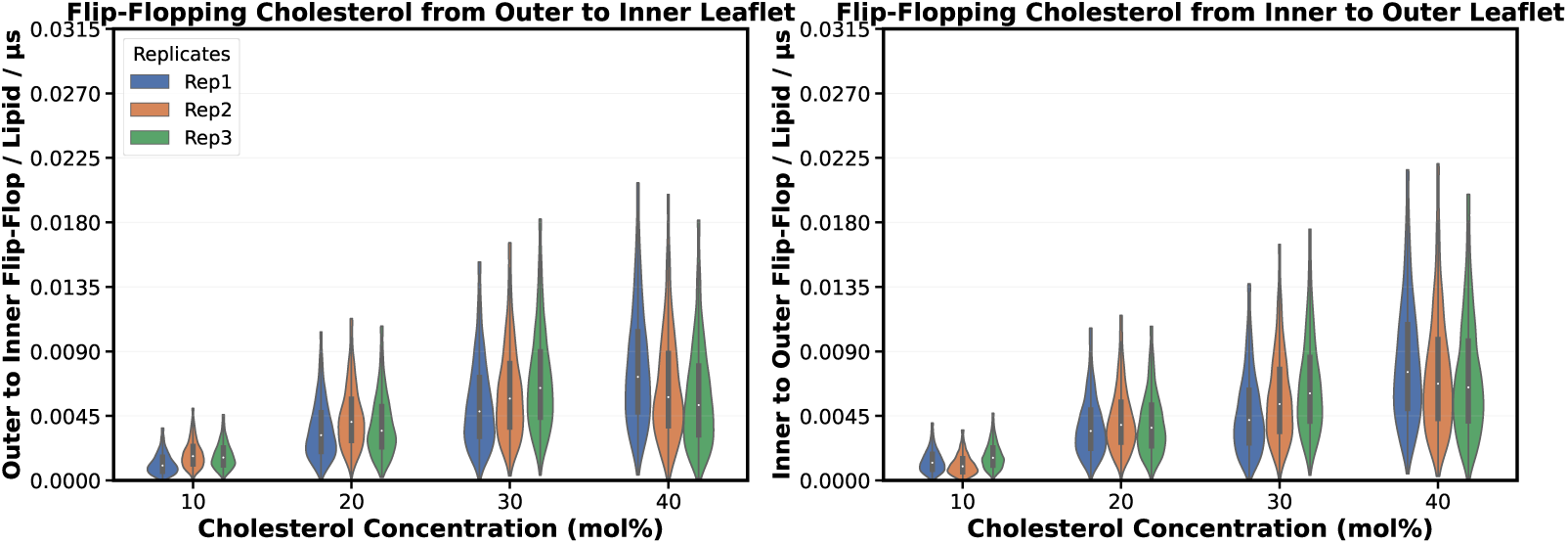
Effect of cholesterol concentration on cholesterol flip-flop in Dry MARTINI vesicles. (Left) Outer-to-inner and (Right) inner-to-outer cholesterol flip-flop frequencies as a function of cholesterol concentration. Both translocation directions exhibit a progressive increase in flip-flop frequency with increasing cholesterol concentration while maintaining approximately balanced exchange between the two membrane leaflets. Each violin plot represents one of three independent 20 µs production simulations.

The monotonic increase in cholesterol flip-flop is consistent with our previous explicit-solvent MARTINI simulations of curved DOPC/cholesterol vesicles [24], indicating that the qualitative dependence on cholesterol concentration is preserved across both solvent representations. The absolute flip-flop frequencies observed in the Dry MARTINI simulations also appear to be higher than those obtained using the explicit-solvent model. This difference is expected because the implicit-solvent representation eliminates explicit water molecules, thereby reducing solvent friction and viscous damping and accelerating molecular motions, including transbilayer migration [14, 41, 52]. Accordingly, the present analysis focuses on the relative influence of cholesterol concentration on flip-flop dynamics rather than on reproducing experimental transbilayer migration rates.

#### 3.6.2. DOPC Flip-Flop

The DOPC flip-flop frequencies are shown in Figure 7. In contrast to cholesterol, DOPC exhibits a non-monotonic dependence on cholesterol concentration in both translocation directions. The flip-flop frequency increases from the cholesterol-free membrane to a pronounced maximum near 20 mol% cholesterol before decreasing at higher cholesterol concentrations.

**Figure 7:**
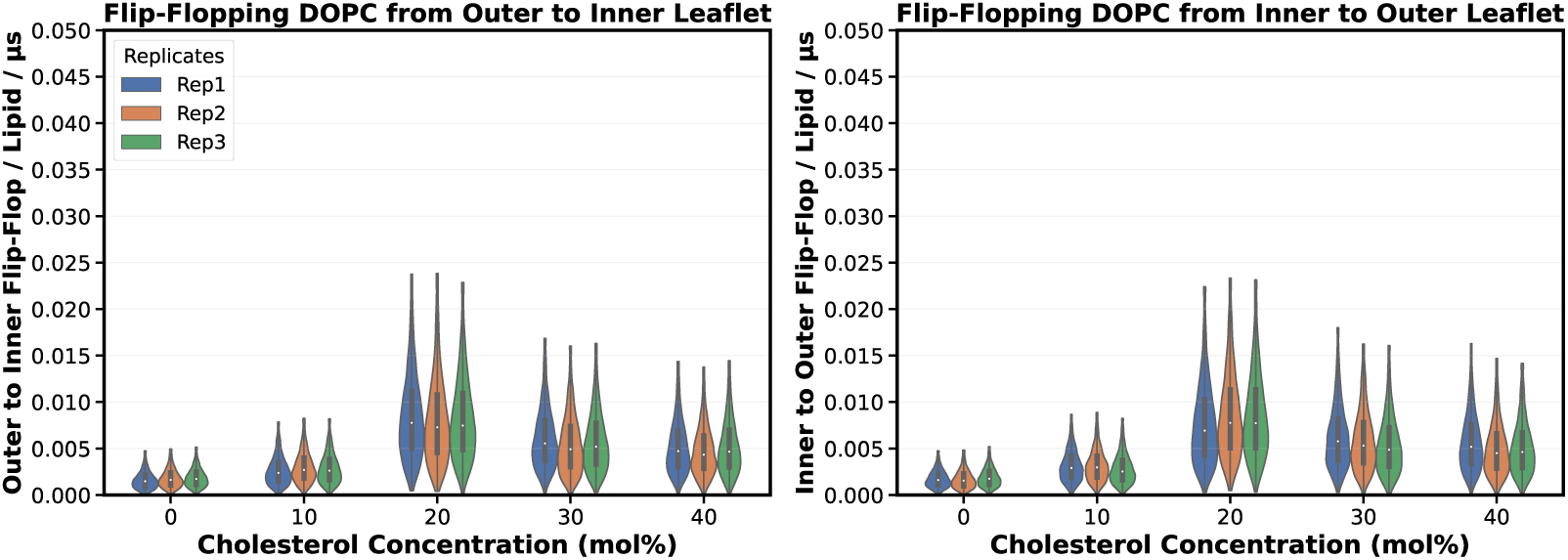
Effect of cholesterol concentration on DOPC flip-flop in Dry MAR-TINI vesicles. (Left) Outer-to-inner and (Right) inner-to-outer DOPC flip-flop frequencies as a function of cholesterol concentration. Both translocation directions exhibit a non-monotonic dependence on cholesterol concentration, with the highest flip-flop frequency occurring near 20 mol% cholesterol, followed by a decrease at higher cholesterol concentrations that nevertheless remains above the cholesterol-free rate through 40 mol% cholesterol. Each violin plot represents one of three independent 20 µs production simulations.

Despite this decline, the flip-flop frequency at 40 mol% cholesterol remains substantially higher than in the cholesterol-free membrane, indicating that cholesterol enhances DOPC transbilayer mobility throughout the concentration range investigated, although the enhancement becomes less pronounced above 20 mol% cholesterol. The outer-to-inner (Figure 7(Left)) and inner-to-outer (Figure 7(Right)) translocation directions exhibit nearly identical concentration dependence, indicating that the observed behavior is independent of the direction of phospholipid translocation.

A similar non-monotonic dependence was observed in our previous explicit-solvent MARTINI simulations of curved DOPC/cholesterol vesicles [24], with the highest DOPC flip-flop frequency occurring near 20 mol% cholesterol. Although the absolute flip-flop frequencies appear to be higher in the Dry MARTINI model than in our previous explicit-solvent simulations, the preservation of this concentration-dependent trend indicates that the response of phospholipid transbilayer migration to cholesterol is robust across both solvent representations. As discussed above for cholesterol flip-flop, the implicit-solvent representation is expected to accelerate transbilayer dynamics relative to explicit-solvent MARTINI.

Above 20 mol% cholesterol, the reduction in DOPC flip-flop coincides with the concentration range in which membrane thickness and lipid-tail ordering increase, while area per lipid, solvent-accessible surface area, and lipid-tail interdigitation all decrease. Together, these structural changes indicate that progressively tighter membrane packing increasingly limits, but does not eliminate, phospholipid transbilayer mobility.

#### 3.6.3. Effect of Cholesterol on Vesicle Shape Anisotropy

The temporal evolution of the relative shape anisotropy (*κ*^2^) of the Dry MARTINI vesicles is shown in Figure 8. All membrane compositions exhibit an initial decrease in *κ*^2^ during approximately the first 3–5 µs of the production simulations, followed by fluctuations around a comparatively stable mean value. This early decrease reflects continued structural relaxation from the initial vesicle configurations, whereas the subsequent plateau suggests that the vesicles reached a stable shape regime with no evidence of large-scale shape transitions during the remainder of the trajectories.

**Figure 8:**
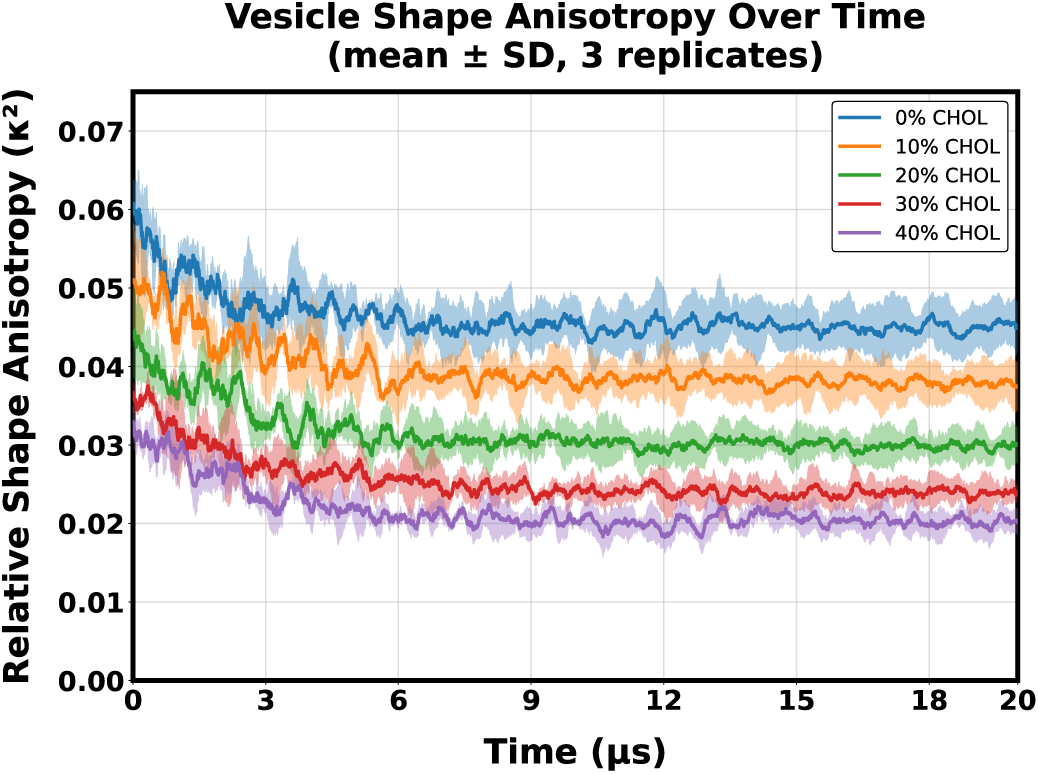
Effect of cholesterol concentration on vesicle shape anisotropy in Dry MARTINI vesicles. Relative shape anisotropy (*κ*^2^) is shown as the mean ± standard deviation across three independent 20 µs simulations for vesicles containing 0–40 mol% cholesterol. All systems exhibit an initial decrease in *κ*^2^ during approximately the first 3–5 µs, followed by fluctuations around a comparatively stable mean value. Increasing cholesterol concentration progressively reduces the average shape anisotropy, indicating that cholesterol-rich vesicles remain closer to spherical symmetry.

Following the initial relaxation period, a clear cholesterol-dependent separation is maintained throughout the simulations, with no crossover between the five membrane compositions. The cholesterol-free vesicle exhibits the highest average relative shape anisotropy and the largest temporal fluctuations, indicating the greatest deviation from spherical symmetry. Increasing cholesterol concentration progressively reduces both the average *κ*^2^ and the magnitude of the associated temporal fluctuations. By 40 mol% cholesterol, the average shape anisotropy is reduced by more than 50% relative to the cholesterol-free vesicle, with the vesicle remaining closest to an isotropic, approximately spherical geometry throughout the simulation.

The reduction in shape anisotropy is consistent with the broader structural changes observed throughout this study. Increasing cholesterol concentration promotes tighter lipid packing, enhanced lipid-tail ordering, increased membrane thickness, reduced interdigitation, and smaller solvent-accessible surface area, all of which could plausibly reduce large-scale vesicle deformations. Previous studies have shown that cholesterol can modify membrane bending rigidity and reduce the amplitude of membrane shape fluctuations in phospholipid membranes [53]. The present results are consistent with this behavior, although we emphasize that *κ*^2^ is a geometric descriptor of vesicle shape and does not by itself provide a direct measure of membrane bending rigidity or other mechanical properties.

The influence of cholesterol on membrane mechanics is known to depend on lipid composition and membrane phase. For example, cholesterol has been reported to increase lipid cooperativity while producing non-universal changes in the mechanical properties of unsaturated membranes [54]. Accordingly, the present results should be interpreted within the context of the simulated DOPC/cholesterol Dry MARTINI vesicles. Within this model, increasing cholesterol progressively reduces global shape anisotropy and the amplitude of vesicle-scale shape fluctuations over the 20 µs simulation period, consistent with enhanced structural stability of the vesicle.

## 4. Conclusion

In this study, we combined the Dry MARTINI force field, the TS2CG membrane builder, and a leaflet-specific restrained-equilibration protocol to investigate how cholesterol influences the structural and dynamical properties of large spherical DOPC vesicles under an implicit-solvent representation. By systematically examining vesicles containing 0–40 mol% cholesterol across three independent 20 µs production simulations for each membrane composition, we established a comprehensive picture of the cholesterol-dependent behavior of curved lipid membranes.

Increasing cholesterol concentration produced a consistent structural reorganization of the membrane. Membrane thickness and lipid-tail ordering increased progressively, whereas the species-specific Voronoi area per lipid, lipid-tail interdigitation, solvent-accessible surface area, and vesicle shape anisotropy all decreased. Together, these complementary structural descriptors indicate that cholesterol drives the membrane toward a more ordered and compact membrane architecture, consistent with its well-established condensing effect in phospholipid bilayers.

Comparison with our previous explicit-solvent MARTINI simulations showed that the principal structural effects of cholesterol are preserved across both Wet and Dry MARTINI models. At the same time, several properties remained sensitive to the solvent representation. Most notably, the species-specific Voronoi packing descriptors exhibited different concentration-dependent behavior in the two models, indicating that local lipid packing is more sensitive to the choice of solvent representation than the overall membrane condensation response. Likewise, lipid-tail interdigitation and the absolute magnitude of lipid flip-flop appeared more pronounced in the Dry MARTINI model. Nevertheless, cholesterol flip-flop increased monotonically and DOPC flip-flop exhibited a reproducible maximum near 20 mol% cholesterol in both models, indicating that the qualitative response of transbilayer lipid mobility to cholesterol is robust across solvent representations.

Overall, these findings demonstrate that Dry MARTINI, combined with an effective restrained-equilibration strategy, provides an efficient and computationally accessible framework for investigating the structure and dynamics of large curved lipid vesicles over microsecond timescales. More importantly, this work demonstrates that comparing explicit- and implicit-solvent coarse-grained models helps identify which membrane properties are preserved across solvent representations and which remain sensitive to the underlying coarse-grained model. Future studies integrating Dry MARTINI simulations with atomistic models and experimental measurements will provide further insight into how cholesterol regulates the structure, dynamics, and organization of complex biological membranes.

## Author Contributions

E.K. conducted simulations, analyzed data, and wrote the manuscript. E.K. assisted with data analysis and molecular dynamics. M.M. designed the research, supervised the project, and reviewed the manuscript.

## Data Availability

All input files and analysis scripts necessary to reproduce the simulations and analyses reported in this study are publicly available at: https://github.com/bslgroup/dryliposo

## Acknowledgments

This research was supported by the NIH (R35GM147423), NSF (CHE 1945465), and the Arkansas Biosciences Institute. Computational resources were provided by the Texas Advanced Computing Center (TACC) at the University of Texas at Austin (Frontera) through LRAC allocation CHE21003. The work also used Stampede at TACC, Bridges-2 at the Pittsburgh Supercomputing Center, and Exapnse at San Diego Supercomputing Center through allocation MCB150129 from the Advanced Cyberinfrastructure Coordination Ecosystem: Services & Support (ACCESS) program. Additional computational support came from the Arkansas High-Performance Computing Center, funded by multiple NSF grants and the Arkansas Economic Development Commission.

