## Supporting Information for "Cholesterol-Dependent Structure and Dynamics of Curved Lipid Vesicles Revealed by Dry MARTINI Simulations"

### 1 Cholesterol Structural Organization

Cholesterol (Fig. S1) was modeled using the MARTINI CG force field [1, 2]. Its amphipathic architecture places the hydroxyl group near the headgroup region while the rigid sterol rings align with lipid acyl chains, allowing cholesterol to modulate packing, increase tail ordering, and enhance membrane mechanical stability [3, 4].

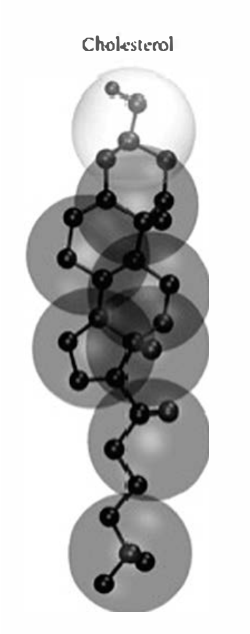


Figure S1: CG representation and structural orientation of cholesterol.

#### 2 CG representation of DOPC

DOPC (Fig. S2) is a zwitterionic phospholipid that forms liquid-disordered, fluid bilayers at physiological temperature due to its unsaturated acyl chains [2, 5, 6]. In the MARTINI representation, DOPC preserves key headgroup polarity and tail flexibility, enabling efficient sampling of sterol–lipid interactions [7].

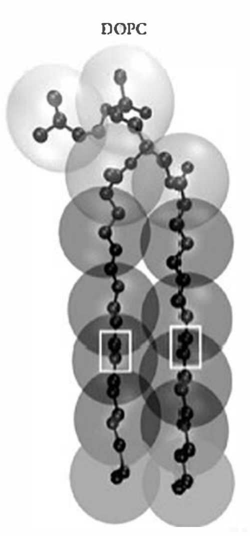


Figure S2: CG representation of DOPC used in the simulations.

### 3 Emergence of Curvature Instabilities During Equilibration

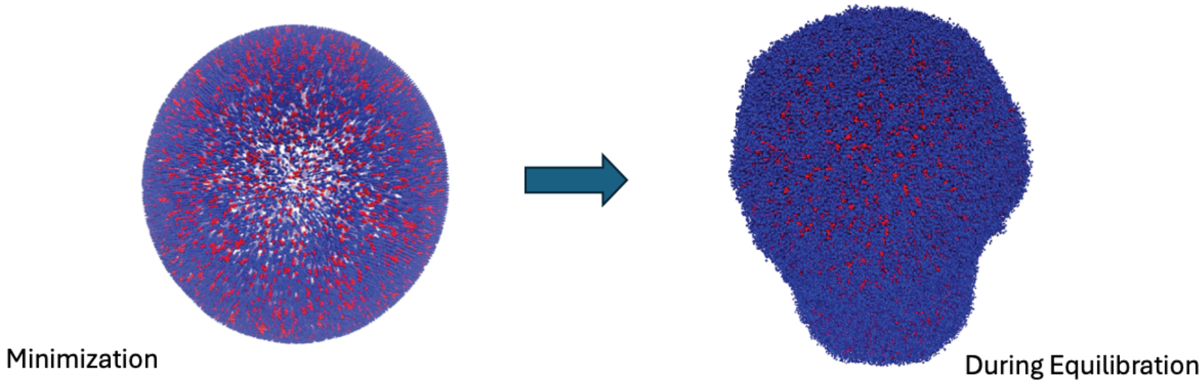


Figure S3: Representative snapshots of a Dry MARTINI DOPC/cholesterol vesicle at 20 mol% cholesterol (310 K): (Left) immediately after energy minimization, showing a smooth, well-defined spherical geometry; (Right) the same vesicle during unrestrained equilibration, showing a pronounced local membrane protrusion (bump). This instability motivated the leaflet-specific, radius-of-gyration-based restrained-equilibration protocol described in Section 2.3.

Although the Dry MARTINI vesicles used in this study were successfully constructed with TS2CG, several challenges emerged during the initial, unrestrained equilibration of these systems. Unlike Wet MARTINI, Dry MARTINI does not include explicit solvent particles, and the solvent-mediated damping that normally stabilizes membrane curvature and suppresses large-scale fluctuations in explicit-solvent simulations is therefore absent. In the absence of this damping, the vesicles developed local membrane protrusions – visible as pronounced surface bumps – during the early stages of unrestrained equilibration.

As shown in Figure S3, a vesicle that is smooth and well-defined immediately after energy minimization can rapidly develop a pronounced local protrusion once unrestrained equilibration begins. This behavior is consistent with the expectation that highly curved membranes are inherently more sensitive to local fluctuations than flat bilayers: the inner and outer leaflets of a vesicle carry different equilibrium radii and packing constraints, and without explicit solvent to damp deviations from the target curvature, these constraints are more easily violated, producing transient protrusions rather than a uniformly curved surface. Cholesterol concentration is expected to further modulate this behavior, since cholesterol alters local membrane rigidity and lipid organization and may therefore shift the balance between curvature-stabilizing and curvature-destabilizing contributions during equilibration.

Because these protrusions reflect transient equilibration artifacts rather than genuine equilibrium membrane behavior, allowing them to develop unchecked would compromise the physical validity of the subsequent production trajectories. This observation directly motivated the leaflet-specific, radius-of-gyration-based restrained-equilibration protocol described in the main text (Section 2.3): by independently restraining the radius of gyration of the inner and outer leaflets during equilibration, the overall vesicle geometry is preserved while local lipid packing is still allowed to relax, preventing the formation of the curvature instabilities illustrated here without imposing any bias on the subsequent, fully unrestrained production simulations.

### 4 Structural Snapshots of Spherical Vesicles

Representative equilibrated configurations of the spherical Dry MARTINI vesicles at 310 K are shown in Figure S4 for the complete cholesterol concentration series (0–40 mol%). These snapshots provide a qualitative visualization of overall vesicle morphology and cholesterol distribution as a function of sterol content, complementing the quantitative structural analyses presented in the main text.

#### 4.1 Spherical vesicles (0–40% cholesterol)

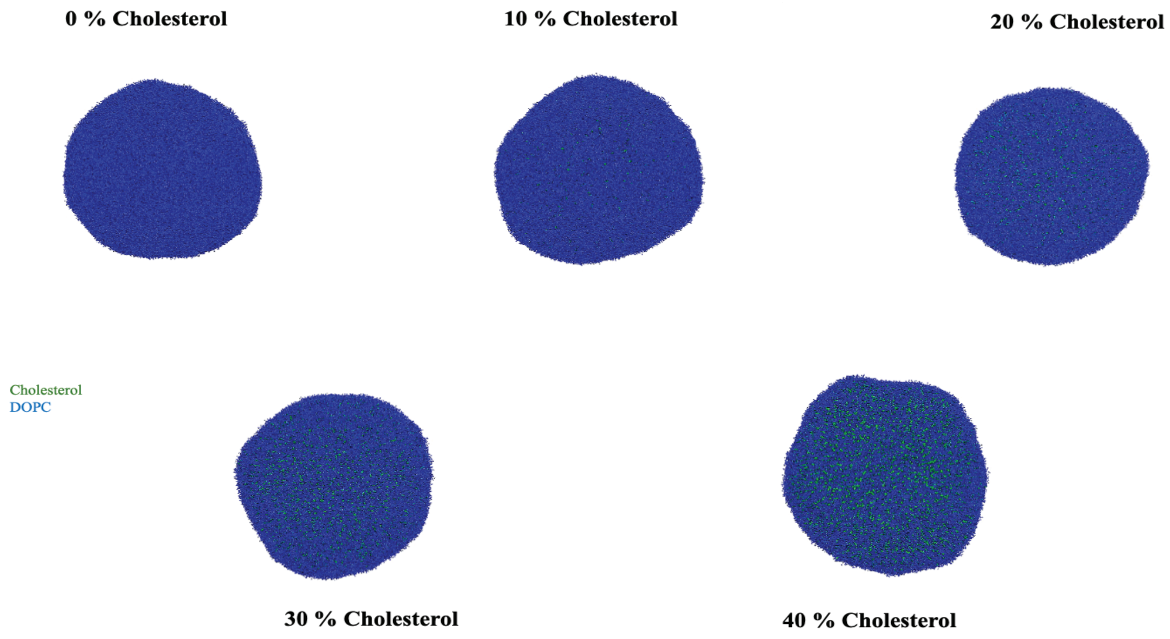


Figure S4: Representative snapshots of spherical DOPC/cholesterol vesicles at 310 K for 0, 10, 20, 30, and 40 mol% cholesterol, with cholesterol symmetrically distributed between the inner and outer leaflets. DOPC and cholesterol molecules are shown in blue and green, respectively. All vesicles retain an overall spherical morphology across the concentration series, consistent with the reduced shape anisotropy quantified as a function of cholesterol concentration in Section 3.6.3, and cholesterol becomes visibly more abundant with increasing sterol content.
